# Drp1-mediated mitochondrial fission protects macrophages from mtDNA/ZBP1-mediated sterile inflammation and inhibits post-infarct cardiac remodeling

**DOI:** 10.1101/2024.10.18.619173

**Authors:** Yuki Kondo, Jun-ichiro Koga, Nasanbadrakh Orkhonselenge, Lixiang Wang, Nao Hasuzawa, Shunsuke Katsuki, Tetsuya Matoba, Yosuke Nishimura, Masatoshi Nomura, Masaharu Kataoka

## Abstract

**Background:** Ischemic heart disease is a leading cause of death worldwide, and heart failure after myocardial infarction (MI) is a growing issue in this aging society. Macrophages play central roles in left ventricular (LV) remodeling after MI. Mitochondria consistently change their morphology, including fission and fusion, but the role of these mitochondrial morphological changes, especially in macrophages, is unknown. This study aims to illuminate the effects and mechanisms of Dynamin-related protein 1 (Drp1), a molecule mediating mitochondrial fission, in macrophages for cardiac remodeling after MI.

**Methods:** This study utilized genetically altered mice lacking Drp1 in monocytes/macrophages (Drp1-KO) to elucidate the specific role of macrophage Drp1 in post-infarct LV remodeling.

**Results:** Deletion of Drp1 in macrophages exacerbated LV remodeling, including reduced ejection fraction and increased LV diameters, which resulted in decreased survival after MI. Histological analysis indicated increased fibrosis and sustained macrophage accumulation in Drp1-KO mice. Blockade of Drp1 in macrophages decreased mitochondrial fission and impaired mitophagy, leading to the subsequent release of mitochondrial DNA (mtDNA) to the cytosol and induction of inflammatory cytokines. This induction was abrogated by an autophagy inducer, Tat-beclin1, or siRNA-mediated knockdown of Z-DNA Binding Protein 1 (ZBP1). Deletion of ZBP1 in bone marrow-derived cells abrogated LV remodeling induced by Drp1 inhibitor, Mdivi-1.

**Conclusion:** Macrophage Drp1 plays a critical role in the pathobiology of LV remodeling after MI, especially mitochondria quality control mechanisms. Macrophage Drp1 could be a novel therapeutic molecule to mitigate the progression of LV remodeling and consequent heart failure after MI.

## Introduction

Ischemic heart disease is the leading cause of death worldwide. In the current era of aging, the incidence of heart failure has been increasing, resulting in the so-called heart failure “pandemic.” Cardiac remodeling following myocardial infarction (MI), particularly of the left ventricle (LV), is still a critical etiology of chronic heart failure despite advancements in early reperfusion therapy for acute MI (AMI). Macrophages, encompassing a heterogeneous population ranging from inflammatory to reparative subsets, play a pivotal role in modulating the healing process after MI. Appropriate inflammation is crucial for tissue repair, which mechanically strengthens the weakened myocardium and prevents severe mechanical complications such as cardiac rupture, whereas excessive inflammation leads to enlargement of LV and inappropriate fibrosis, resulting in systolic and diastolic dysfunction (*1*). However, the regulatory mechanisms of macrophage function and polarization during the healing process after MI remain largely unexplored.

We have recently elucidated the role of dynamin-related protein 1 (Drp1), a molecule that promotes mitochondrial fission, in macrophage functions both in vivo and in vitro. In a mouse vascular injury model, deletion of macrophage Drp1 attenuated inflammation and intimal thickening after mechanical injury of femoral arteries. In ex vivo cultured macrophages, blockade of Drp1 inhibited the production of mitochondria-derived reactive oxygen species (ROS) and expression of molecules associated with inflammation (*2*). We have previously reported that the Drp1 inhibitor, Mdivi-1, decreased infarct size after myocardial ischemia-reperfusion through the inhibition of mitochondrial outer membrane permeabilization (*3*).

Cardiomyocyte-specific deletion of Drp1 increased infarct size after ischemia-reperfusion in the heart (*4*) and exacerbated heart failure after pressure overload of LV by transverse aortic constriction (*5*).

Drp1 is also involved in the pathobiology of various diseases in other organs. In the liver, hepatocyte-specific deletion of Drp1 exacerbated LPS-induced acute liver injury by promoting ER stress response and inflammasome activation (*6*). In the kidney, the deletion of Drp1 protected the podocytes from albuminuria-associated damage in diabetes (*7*). These findings indicate that the role of Drp1 varies depending on the disease and the cellular context. It is, however, unknown whether mitochondrial fission and fusion, called mitochondrial dynamics, is a regulator of macrophage function and polarization in the healing process after MI. Therefore, this study aims to elucidate the role of Drp1-mediated mitochondrial fission in macrophage-mediated inflammation and LV remodeling after MI.

## Methods

### Animal studies

The Ethics Committees approved all the experimental protocols for Animal Care and Experimentation at the University of Occupational and Environmental Health and Kyushu University. All the animal care and experimental procedures were conducted in strict accordance with the NIH Guide for the Care and Use of Laboratory Animals. All animal surgeries were performed under inhaled isoflurane anesthesia to minimize animal suffering. We generated floxed Drp1 mice on a C57Bl/6J background, as previously described (*2*), and crossed these with lysozyme M Cre mice on a C57Bl/6J background. Drp1^flox/flox^; LysM^Cre/Cre^ genotype was designated as Drp1-KO (Drp1 knockout), while littermates with the Drp1^+/+^; LysM^Cre/Cre^ genotype served as controls. ZBP1-KO mice were developed using frozen embryos supplied by the Laboratory Animal Resource Bank at the National Institutes of Biomedical Innovation, Health, and Nutrition in Japan (Osaka, Japan). The first-generation heterozygous offspring resulting were subsequently interbred to produce homozygous ZBP1-KO mice.

### Myocardial infarction by ligation of left anterior descending coronary arteries in mice

Mice aged 8 to 12 weeks underwent surgery to induce MI, as described previously (*1*). The procedure involved anesthetizing the mice with isoflurane, tracheal intubation, and a lateral thoracotomy at the third intercostal space. Once the epicardium was exposed, the left anterior descending (LAD) coronary artery was located and ligated approximately 2 to 3 millimeters below the left atrium using an 8-0 nylon suture. The surgical incisions were then closed with a 5-0 silk suture. The Drp1 inhibitor Mdivi-1 (Sigma-Aldrich, Burlington, Massachusetts) was dissolved in dimethyl sulfoxide (DMSO) and intraperitoneally injected using Alzet^Ⓡ^ mini-osmotic pumps (model 2004, Charles River, Wilmington, MA) at a dosage of 50 mg/kg in mice indicated elsewhere.

### Echocardiography

Transthoracic echocardiography was conducted using an 18 MHz linear probe (GE L8-18i-SC) and a GE Vivid E9 Ultrasound Machine (GE Healthcare Japan, Tokyo, Japan). The mice were placed in a supine position and anesthetized with isoflurane, maintaining anesthesia at a level that kept the heart rate at approximately 500 beats per minute. LV end-diastolic diameter (LVEDD), LV end-systolic diameter (LVESD), interventricular septal wall thickness (IVSWT), LV posterior wall thickness (LVPWT), and LV ejection fraction (LVEF) were measured by M-mode image at the level of the papillary muscle. All echocardiographic analyses were conducted by investigators blinded to the treatment groups.

### Histopathology and Immunohistochemistry

For histological analyses, frozen sections of the heart were prepared as previously described (*1*). Briefly, the mice were euthanized by intravenous administration of 100 mg/kg pentobarbital. Blood was washed by injection of PBS from the LV apex, and perfusion fixed with 10% neutral buffered formalin. The hearts were excited and fixed again for overnight. Then, the hearts were sequentially incubated in 15% and 30% sucrose solutions for 24 hours each and embedded in an OCT compound (Sakura Finetek, Tokyo, Japan). Sections were cut using microtome with 9 μm thickness. These sections were stained using Masson’s trichrome (ScyTek Laboratories, Logan, UT) according to the manufacturer’s instructions. Immunostainings were performed using the following antibodies: rat anti-mouse Mac-3 monoclonal antibodies (550292; BD Pharmingen, San Diego, CA)and Cleaved Caspase-3 rabbit monoclonal antibodies (9664; Cell Signaling Technology, Danvers, MA). Following incubation with peroxidase polymer-modified secondary antibodies (MP-7451; Vector Laboratories, Newark, CA), 3-amino-9-ethylcarbazole (Vector Laboratories, Inc.) was used as a substrate for visualization. Sections were counterstained with Mayer’s hematoxylin, and microscopic observations were performed using the ECLIPSE Ci-L plus microscope equipped with a digital image analysis system (Nikon, Kanagawa, Japan) and NIS-Elements software. In immunofluorescence staining, sections were incubated with Alexa Fluor-conjugated secondary antibodies and imaged using a confocal laser scanning microscope (LSM 880; Carl Zeiss AG, Oberkochen, Germany).

### Flow cytometry

Excised hearts were enzymatically digested using a mixture containing 450 U/mL collagenase type I (17100017; Thermo Fisher Scientific, Waltham, MA), 125 U/mL collagenase type XI (C9407; Sigma-Aldrich), 60 U/mL DNase I (18047019; Sigma-Aldrich), and 60 U/mL hyaluronidase (H3506; Sigma-Aldrich) in PBS with 20 mM HEPES, maintained at 37 °C for one hour. After digestion, cell suspensions were centrifuged at 240 x g for 10 minutes at 4 °C. Then, the supernatants were discarded, and pellets were resuspended in 50 μl of HBSS (14025092; Thermo Fisher Scientific) with 1% FBS. Fc receptors were blocked using mouse seroblock FcR (BUF041A; Bio-Rad Laboratories, Hercules, CA), followed by incubation with a cocktail of monoclonal antibodies including CD45-APC, CD90.2-PE, B220-PE, CD49b-PE, Ly-6G-PE, CD11b-APC-Cy7, and Ly-6C-FITC (BD Pharmingen) for one hour at 4 °C. To exclude nonviable cells, 7-AAD was added 20 minutes before flow cytometry, and analyses were performed using a CytoFLEX S flow cytometer (Beckman Coulter, Inc., Indianapolis, Indiana). Isotype IgG (BD Pharmingen) was used to stain control samples. Monocytes/macrophages were identified as CD45^+^CD11b^+^ Lineage (CD90/B220/CD49b/NK1.1/Ly-6G)(Lin)^-^ Ly-6C^high/low^, the neutrophils were identified as CD45^+^CD11b^+^ Lin^+^, and the lymphocytes were identified as CD45^+^CD11b^-^Lin^+^.

### Transmission electron microscopy

To prevent cardiac tissue desiccation, normal saline was continuously applied. Infarcted areas were excised into one-millimeter cubes and fixed in a mixture of 2% glutaraldehyde and 2% paraformaldehyde in 0.1M phosphate buffer. After pre-fixation, samples were washed and post-fixed in 1% osmium tetroxide in phosphate buffer for 1 to 2 hours, then dehydrated through a graded ethanol series from 50% to 100%. Following dehydration, samples were infiltrated with synthetic resin, starting with a QY-1-to-resin ratio of 1:2, progressing to 1:1, and culminating in 100% resin, each stage lasting 20 to 30 minutes. The fully infiltrated samples were embedded in resin within beem capsules and polymerized at 60°C for three days. Ultrathin sections (80 nm) were cut and stained first with 4% uranyl acetate and then with Reynold’s lead citrate solution prepared from lead nitrate, sodium citrate, distilled water, and sodium hydroxide.

### Preparation of mouse peritoneal macrophages

Three days prior to collection, 2 ml of 3% Brewer thioglycolate medium was injected into the peritoneal cavity to elicit macrophages. On the collection day, following euthanasia by cervical dislocation, a sterile midline incision was made carefully to avoid damaging the peritoneum. Subsequently, 5 ml of PBS was injected into the peritoneal cavity, and the carcass was gently agitated to facilitate the distribution of PBS. The peritoneal fluid, now containing the macrophages, was then aspirated. Macrophage counts were performed using the Countess 3FL (Thermo Fisher Scientific), and cells at a concentration of 1.0×10^6^ cells/ml were cultured in 5-7 ml of RPMI-1640 medium (RNBK4492: Sigma-Aldrich) supplemented with 10% fetal bovine serum (12003C-500ML; Sigma-Aldrich) and 1% penicillin. The macrophages were then incubated at 37°C and analyzed 24 hours later.

### Cell cultures

RAW264.7 cells (TIB-71: ATCC, Manassas, VA) were maintained in Dulbecco’s Modified Eagle Medium (DMEM) supplemented with L-Glutamine (048-30275; FUJIFILM Wako Pure Chemical Corporation), 10% fetal bovine serum and 1% penicillin. These cells were treated with 10 ng/mL LPS (L5293; Sigma-Aldrich) and 10 ng/mL mouse IFN-γ (11276905001; Roche, Basel, Switzerland) to induce polarization toward inflammatory macrophages. Mdivi-1 (50 μmol/L) dissolved in DMSO was used to inhibit Drp1 to inhibit Drp1 in these macrophages. Tat-beclin 1 D11 Autophagy Inducing Peptide 1 (NBP2-49888; Bio-Techne Corporation, Minneapolis, MN) was used to induce autophagy and VBIT-4. To culture cells under hypoxic conditions with 0.5% O2, the BIONIX-1 hypoxic culture kit (Sugiyamagen, Tokyo, Japan) was used according to the manufacturer’s instructions.

### Visualization of mitochondria in in vitro cultured macrophages

For visualization of mitochondria, cells were stained with MitoTracker™ DEEP Red (M22426; Thermo Fisher Scientific), a mitochondrion-selective fluorescent probe (Ex/Em=644/655 nm). Stained macrophages were mounted on slide glasses and examined under a BZ-X800 Fluorescence microscope (Keyence, Osaka, Japan). The assessment of mitochondrial fission was conducted following previously established methods (*14*). Briefly, mitochondria fragmentation count (MFC) was calculated by counting non-contiguous segments of mitochondria and dividing the result by the total pixel count that forms the mitochondrial network. A higher MFC indicates a greater degree of mitochondrial division.

### In vitro loss-of-function experiments

siRNA-mediated knockdown was performed to investigate the role of Drp1 and ZBP1. RAW264.7 cells were transfected with 20 nM of either ON-TARGETplus mouse Drp1 siRNA (L-054815-01-0005), ON-TARGETplus mouse Zbp1 siRNA (L-048021-00-0005), or non-targeting siRNA (D-001810-10-20) (all from Horizon Discovery, Cambridge, UK), using PolyMag Neo (PG61000; OZ Biosciences, Marseille, France) following the manufacturer’s instructions. To induce mitophagy, cells were treated with 1 μg/ml oligomycin (75351; Sigma-Aldrich) and 30 μM CCCP (215911; Sigma-Aldrich) for 1 hour starting from 36 hours after siRNA transfection.

### Mitochondrial Isolation and Western blotting

Mitochondria were isolated from cultured cells at specified time points using the Mitochondria Isolation Kit for Cultured Cells (89874; Thermo Fisher Scientific). For protein analysis, mitochondria were lysed with M-PERTM (WL337237; Thermo Fisher Scientific) to obtain samples containing 10-30 μg of protein quantified by BCA Protein Assay Kit (XD340639; Thermo Fisher Scientific). Immunoblottings were performed with the following primary antibodies after electroporation: against VDAC (4661S), LC3 A/B (12741S), β-actin (A3854), AMPKα (2532S), Phospho-AMPKα (Thr172) (2531S), DRP1 (8570S), Phospho-DRP1 (Ser616) (4494S), Phospho-PINK1 (Ser228) (Cell Signaling Technology), STAT2 (MA5-42463), Phospho-STAT2 (Thr387) (PA5-114647) and PINK1 (PA1-16604) (Thermo Fisher Scientific). Blots were incubated with HRP-conjugated rabbit IgG (NA934-1ML; Amersham plc), developed with AmershamTM ECL reagent (RPN2236; GE Healthcare). Images were captured with the Amersham™ ImageQuant™ 500 system (Cytiva), and the intensities of the band were quantified using ImageQuant TL software (Cytiva).

### Observation of mitophagy in RAW267.4 cells

mito-Keima Red (pMitophagy Keima-Red mPark2) (Ex/Em=550/610 nm) (AM-V0259M; BML, Tokyo, Japan), a pH-dependent fluorescent protein with mitochondria targeting sequence, was employed to assess mitophagy. mito-Keima was transfected to target cells using PolyMag Neo (PG61000; OZ Biosciences) according to the instructions. Mitophasic cells were observed by laser scanning confocal microscopy (LSM880; Carl Zeiss) and quantified by a fluorescent plate reader (SH-1000; CORONA ELECTRIC Co., Ibaraki, Japan)) using a wavelength of 586 nm.

### Extraction of cytosolic mitochondrial DNA

Cytosolic fractions of RAW264.7 cells without mitochondria were prepared by the Mitochondria Isolation Kit for Cultured Cells (ThermoFisher Scientific), and DNA was extracted using the DNeasy Blood & Tissue Kit (QIAGEN, Venlo, Netherlands). Semi-quantitative real-time PCR was performed to amplify mtDNA-encoded cytochrome c oxidase I genes.

### Semi-quantitative real-time PCR

RNA was extracted from cells using the ReliaPrep™ RNA Cell Miniprep System (Promega, Madison, WI). Subsequent cDNA synthesis was conducted using PrimeScript RT Master Mix (RR036A; Takara Bio, Shiga, Japan). Real-time PCR analyses were performed on a QuantStudio™ 3 Real-Time PCR System (Applied Biosystems™, Waltham, MA) using PowerTrack™ SYBR™ Green Master Mix (01339232; Thermo Fisher Scientific), with specific primers detailed in Supplemental Table 1. The expression data were calculated using the ΔΔCT method and normalized by internal controls according to the experimental conditions.

### Transcriptome analysis of cultured macrophages

Total cellular RNA was extracted using ISOGEN (311-02501; NIPPON GENE, Tokyo, Japan) from RAW267.4 cells at 24 or 96 hours after treatment with 10 ng/mL LPS and 10 ng/mL mouse IFN-γ, with or without 50 µM Mdivi-1. Subsequent next-generation RNA sequencing was outsourced and conducted at Kyushu Pro Search LLP (Fukuoka, Japan) using Nextseq 500 (Illumina, San Diego, CA). Lead count data were analyzed using integrated Differential Expression & Pathway Analysis 2.01 (iDEP2.01) (*23*) and Gene Set Enrichment Analysis (GSEA) software v4.3.3 (*23–24*). Differentially expressed genes (DEGs) were identified by the limma-voom method with a fold change of >2 or <0.5. Gene Ontology (GO) analysis was performed using Metascapse (*25*), and enriched pathways were identified as gene sets with adjusted p-value<0.05. Network analyses of DEGs were performed STRING (*26*). Cytoscape version 3.10.2 (*27*) and cytoHubba 0.1. (*28*) were used to identify Hub genes.

### Detection of Apoptotic cells

Apoptotic cells were stained using an In situ Apoptosis Detection Kit (MK500; Takara Bio). Briefly, cells were fixed with 4% paraformaldehyde, followed by washes with PBS. Cells were then permeabilized by 100 μl of permeabilization buffer and incubated with a mixture of TdT enzyme and labeling buffer in a humidity chamber. Apoptotic cells emitting fluorescences were observed using an LSM880 microscope with a 488 nm laser (Carl Zeiss).

### Bone marrow transplantation

Bone marrow transplantation was performed as previously reported (*29*). Donor mice aged 6–8 weeks were euthanized via cervical dislocation, and bone marrow cells (BMCs) were harvested from the femurs, tibias, and humeri of the donor mice. Approximately 4×10^7^ unfractionated BMCs were collected from each mouse, and 5×10^6^ cells suspended in PBS were transplanted into each recipient mouse via the tail vein. Age-matched recipient mice were irradiated with a total of 7 Gy X-rays on the day before the injection of BMCs. Three weeks after the BMC transplantation, the mice were used for experiments. In this way, myeloid ZBP1-KO mice were generated.

### Statistical analyses

Data are presented as means ± SD unless specified otherwise. Statistical differences between the two groups were analyzed using an unpaired t-test. For three or more groups, differences were assessed using one-way or two-way ANOVA, followed by Tukey’s post-hoc multiple comparison test, performed with Prism Software version 9.0 (GraphPad Software, San Diego, CA). A p-value of less than 0.05 was considered statistically significant.

## Results

### Deletion of Drp1 in Lys M-positive cells accelerated LV remodeling after MI

To clarify the selective role of macrophage Drp1 in the mechanisms of LV remodeling after MI, we have prepared genetically altered mice by crossing floxed Drp1 mice and lysozyme M (Lys-M) Cre mice (Drp1-KO). There were no significant differences in hemodynamic parameters, including heart rate and systolic blood pressure measured by tail-cuff method between control Lys-M Cre (control) and Drp1-KO mice (Fig.1b, c). Echocardiography on 3, 7, 14, and 28 days after LAD ligation (Fig.1d-i) demonstrated a significant reduction of LV ejection fraction (LVEF) in Drp1-KO mice compared to control mice, which was observed from 14 days after MI and progressed until 28 days (Fig.1e). Deletion of macrophage Drp1 increased end-diastolic and end-systolic LV diameters determined by M-mode measurement of echocardiography, which was observed from 7 days after MI and progressed until 28 days (Fig.1f, g). Interventricular septum and posterior wall thickness were comparable between the control and Drp1-KO group after MI (Fig.1h, i). As shown in the Kaplan-Meier curve in Fig.1j, deletion of macrophage Drp1 decreased survival after MI as compared to control mice (25.00 vs. 63.64%, p=0.002) (Fig.1j).

**Fig. 1.**
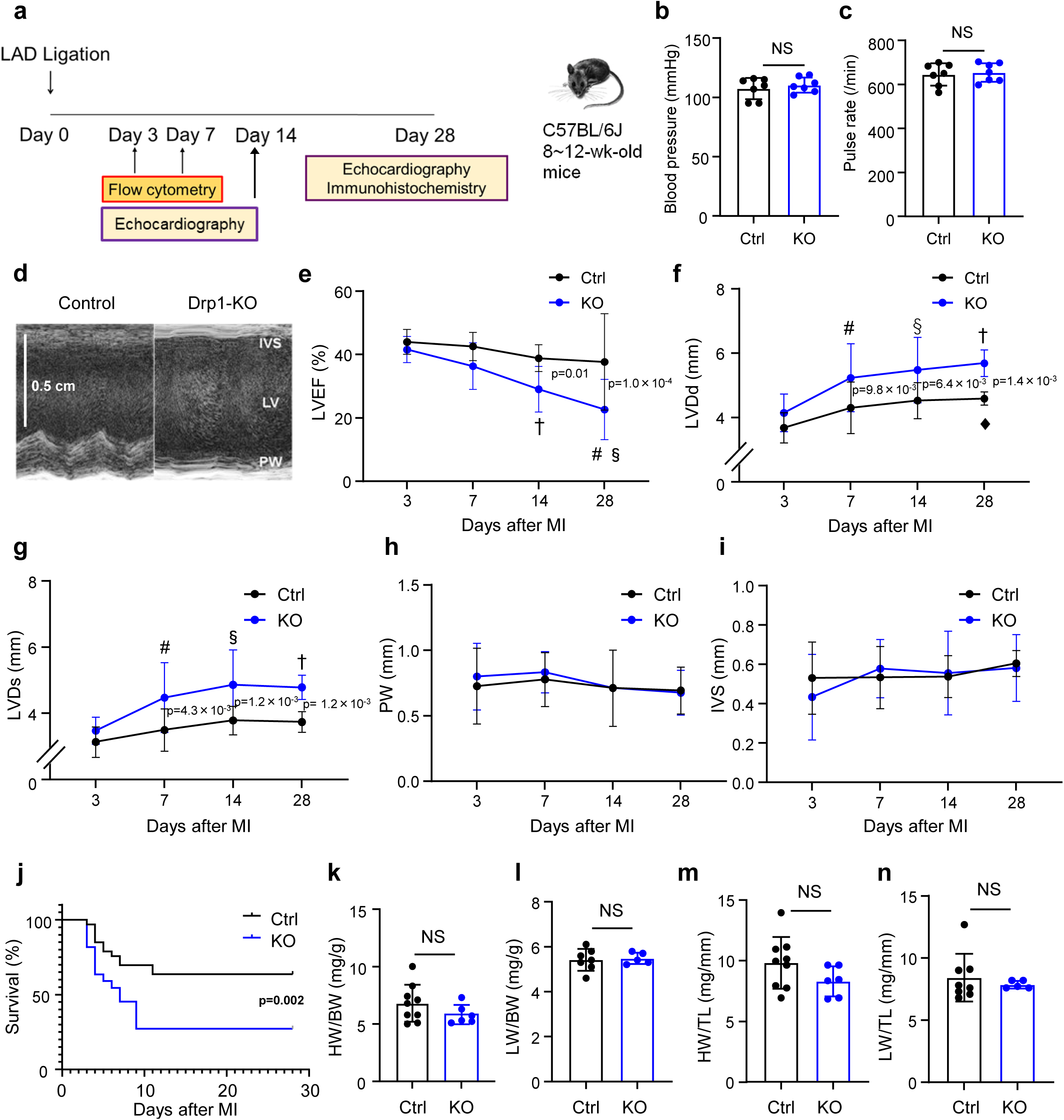
Deletion of Drp1 in Lys M-positive cells impairs systolic function and promotes LV dilatation after MI. (**a**) Experimental protocol. LAD was ligated at 8 to 12 weeks of age in lysozyme-M^Cre/Cre^Drp1^+/+^ (Control) and lysozyme-M^Cre/Cre^Drp1^flox/flox^ (Drp1-KO) mice. (**b-c**) Hemodynamic parameters of mice on day 28 (blood pressure; p=0.49, pulse rate; p=0.73, unpaired t-test) (N=7, each). (**d-i)** Time course of LV function and morphology measured by echocardiogram. (**e**; #p<0.0001 vs. Day 3, §p=0.0027 vs. Day 7, †p=0.0067 vs. Day 3, **f**; #p=0.008 vs. Day 3, §p=0.0008 vs. Day 3, †p= <0.0001 vs. Day 3, p=0.035 vs. Day 3, **g**; #p=0.01 vs. day 3, §p=0.0002 vs. day 3, †p=0.0004 vs. day 3, **h**; there was no significant difference between groups, **i**; there was no significant difference between groups Two-way ANOVA) (N=8-10). (**j**) Kaplan-Meier curve showing survival after LAD ligation (Log-rank test) (N=30 and 33). (**k-n**) Weight of the heart and lung measured at 28 days after MI (**k**; p=0.19, **l**; p=0.79, **m**; p=0.14, **n**; p=0.53 unpaired t-test) (N=5-9). All data are presented as mean ± s.e.m. Ctrl: control, KO: Drp1-KO, HW: heart weight, BW: body weight, LW: lung weight, TL: tibia length. NS: not significant.

**Table 1.**
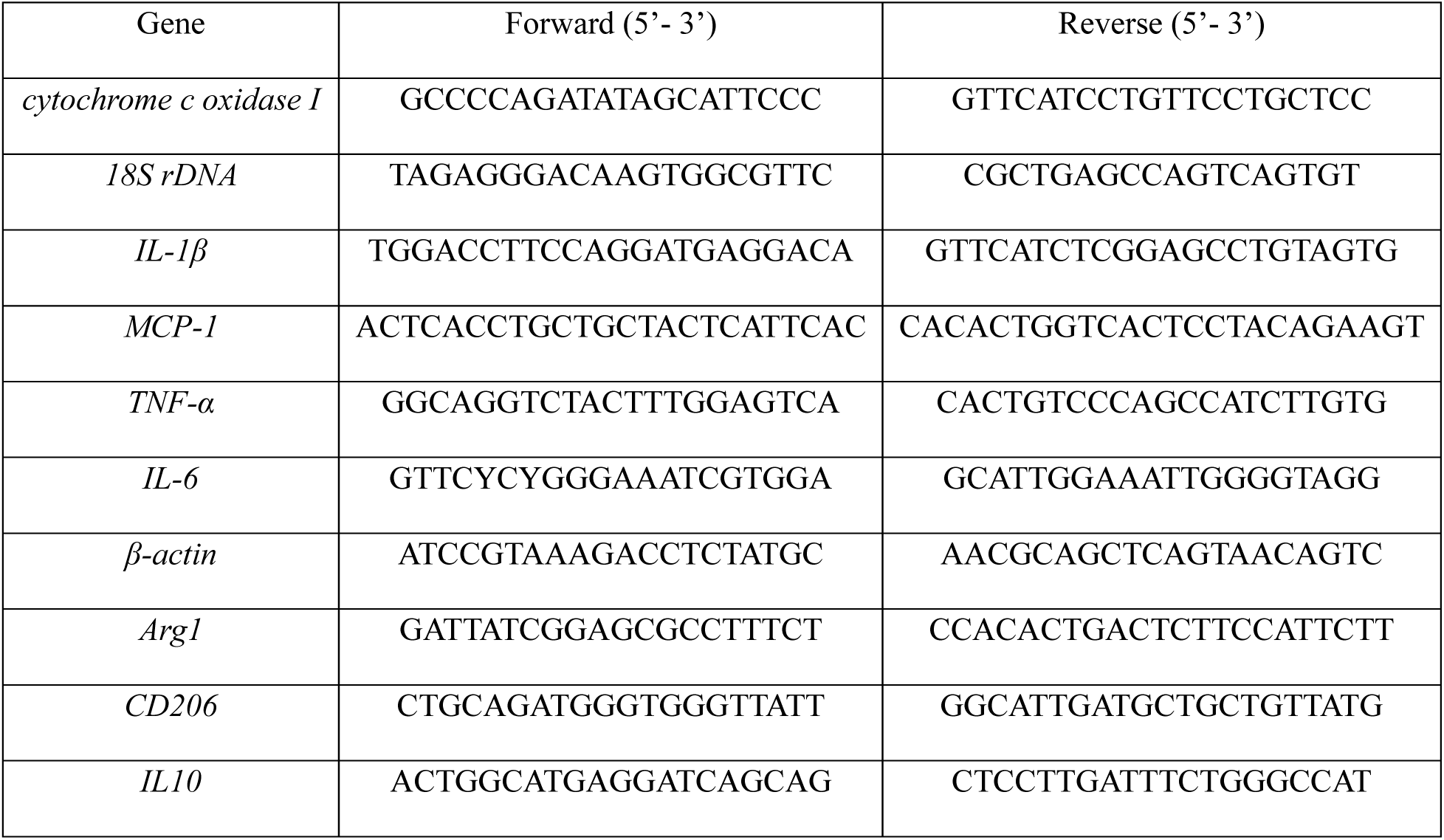
Primer pairs used in this study.

Especially, deaths in both groups have occurred until 10 days after MI. Measurements of the heart and lung weights at the endpoint of the study showed no significant differences between Drp1-KO and control mice (Fig.1k-n).

### Deletion of Drp1 in Lys M-positive cells increased LV fibrosis and caused sustained inflammation after MI

Masson’s trichrome staining revealed that the fibrosis in the infarcted heart was increased by the deletion of macrophage Drp1 as compared to the control (9.78 vs. 6.57%, p=0.018) (Fig.2a). In addition, the deletion of Drp1 significantly increased the circumferential length of the infarcted area (45.94 vs. 26.96%, p= 2.0×10^-4^). In contrast, there were no significant differences in compensated cardiac hypertrophy in the non-infarcted area quantified as the cross-sectional area of cardiomyocytes (Fig.2b, c).

**Fig. 2.**
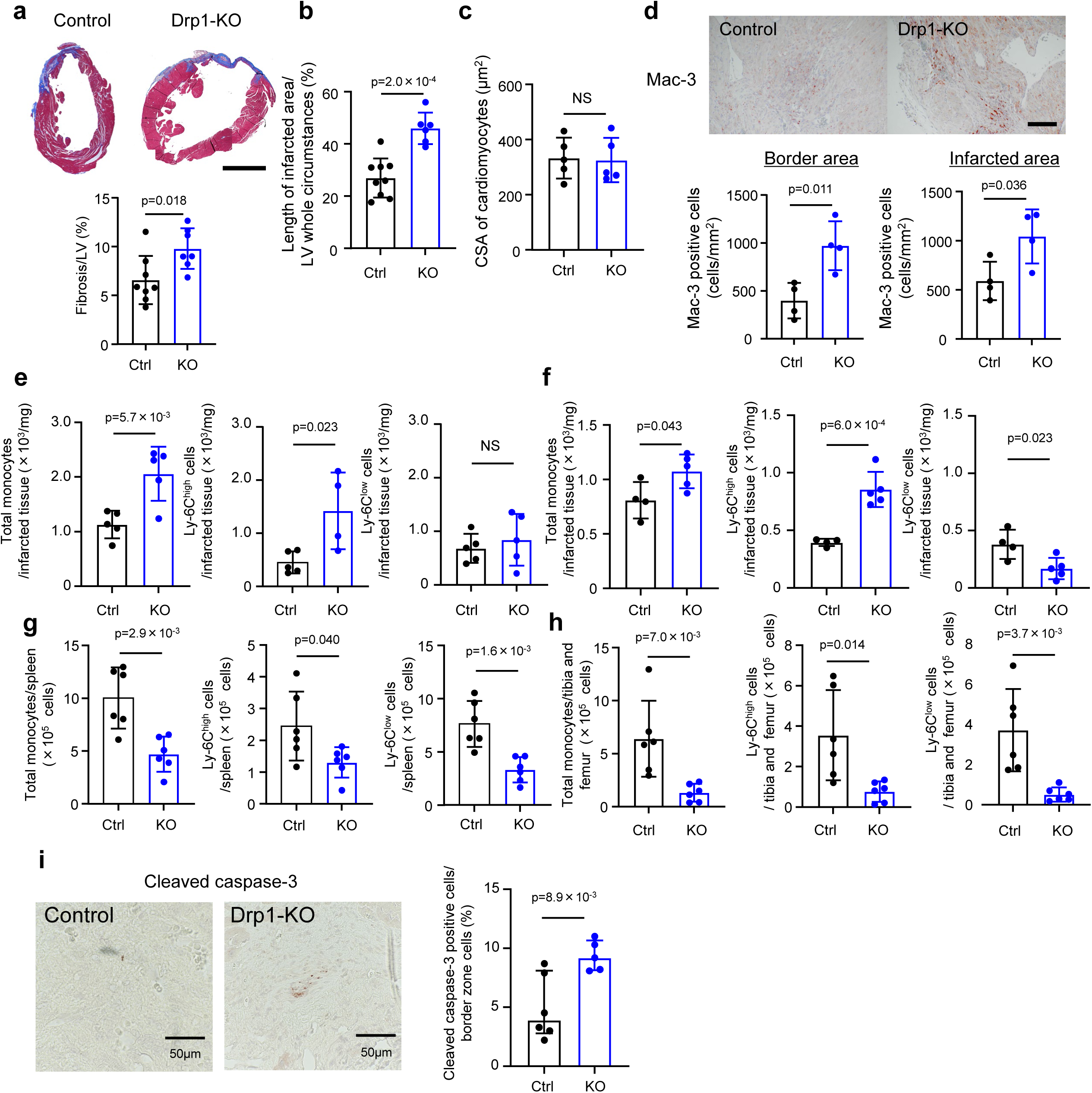
Deletion of Drp1 in Lys M-positive cells exacerbates LV remodeling after MI and causes sustained macrophage accumulation. (**a**) Masson’s trichrome staining of the infarcted heart. Fibrosis was quantified as the ratio of fibrotic area to the total LV area (unpaired t-test) (N=7-8). The scale bar indicates 2 mm. (**b**) The relative length of scar area to whole circumstances (unpaired t-test) (N=6-9). (**c**) Cross-sectional area of cardiomyocytes in non-infarcted area (p=0.89 unpaired t test) (N=5). (**d**) The number of mac-3 positive cells in the border zone and infarcted area (unpaired t-test) (N=4). The scale bar indicates 50 μm. (**e, f**) Flow cytometry analysis showing the number of macrophages accumulating infarcted hearts on Day 3 (e) and Day 7 (f) (Ly-6C^low^ cells on Day 3; p=0.54 unpaired t-test) (N=4-5). (**g, h**) Flow cytometry analysis showing the number of macrophages in the spleen (g) and bone marrow (h) on Day 3 (unpaired t-test) (N=6). (**i**) Immunostaining of cleaved caspase-3 in the infarcted heart on Day 28. The right graph shows quantitative results in the periinfarct border zone. (unpaired t-test) (N=5-6). All data are presented as mean ± s.e.m. Ctrl: control, KO: Drp1-KO.

Immunostaining of Mac-3 at 28 days after MI indicated increased accumulation of macrophages in the infarct area and peri-infarct border zone, in either of which, macrophage accumulation was enhanced by the deletion of macrophage Drp1 (infarct area: 1043.0 vs. 588.6 cells/mm^2^, p=0.011, border zone: 971.4 vs. 397.8 cells/mm^2^, p=0.036) (Fig.2d). Moreover, flow cytometry showed that numbers of either total and Ly-6C^high^ inflammatory monocytes in the infarct area were increased in the Drp1-KO mice compared to the control mice at day 3 after MI (total monocytes: 2060 vs 1129/mg, p=5.7×10^-3^, Ly-6C^high^ monocytes: 1421 vs 449/mg, p=0.023) (Fig.2e). The gating and histograms of the flow cytometry are shown in Extended Data Fig.1. As shown in the histogram, the Drp1-KO mice have a higher number of Ly-6C^high^ monocytes with a unimodal distribution on day 3 after MI, while the control mice have a bimodal distribution of Ly-6C^high^ and Ly-6C^low^ monocytes. In contrast, on day 7 after MI, the control mice exhibit an unimodal distribution of Ly-6C^low^ monocytes, whereas the Drp1-KO mice still show an unimodal distribution with a higher number of Ly-6C^high^ monocytes. On day 7 after MI, increased numbers of total and Ly-6C^high^ monocytes were still observed in Drp1-KO mice (856.1 vs 394.7/mg, p=6.0×10^-4^). In contrast, the Ly-6C^low^/CD206^+^ monocytes/macrophages were decreased compared to the control group (170.5 vs 380.6/mg, p=0.023) (Fig.2f). A reduction in total monocytes/macrophages, Ly-6C^high^ monocytes, and Ly-6C^low^ monocytes/macrophages in the spleens and bone marrows were observed in the Drp1-KO mice at day 3 after MI (Fig.2g, h).

Hence, we have assessed the extent of apoptosis in the infarcted heart to address the possibility that decreased apoptosis in Drp1-KO mice caused increased macrophage accumulation. Immunohistochemical analysis of cleaved caspase-3 at day 28 after MI indicated that deletion of macrophage Drp1 rather increased the ratio of cleaved caspase-3^+^ cells in the peri-infarct border area (9.36 vs 4.93%, p=8.9×10^-3^) (Fig.2i).

### Hypoxia and starvation induced AMPK phosphorylation and Drp1-mediated mitochondrial fission

Macrophages accumulating in the infarcted area are exposed to hypoxia and loss of energy sources due to disruption of blood supply. Therefore, we have examined whether hypoxia and starvation induce activation of Drp1 and subsequent mitochondrial fission in peritoneal macrophages. Skewing to inflammatory macrophages induced by lipopolysaccharide (LPS) and interferon-γ (IFN-γ), increased adenosine monophosphate-activated protein kinase (AMPK), a master regulator of cellular metabolism, and its phosphorylation. In contrast, the ratio of p-AMPK/AMPK was not increased by LPS and IFN-γ (Fig.3a). Signal transducer and activator of transcription 2 (Stat2) and PTEN-induced putative kinase 1 (PINK1), which have been reported as activators of Drp1, were also evaluated (Extended Data Fig.2). In these macrophages, a combination of hypoxia and starvation increased p-AMPK/AMPK (Fig.3b). Western blot demonstrated LPS and IFN-γ induced Drp1 expression and its phosphorylation at Ser616, which was further enhanced by hypoxia and starvation. The p-Drp1(Ser616)/Drp1 ratio was increased only in conditions with hypoxia and starvation, which was mitigated by an AMPK inhibitor (Fig.3c, d).

**Fig. 3.**
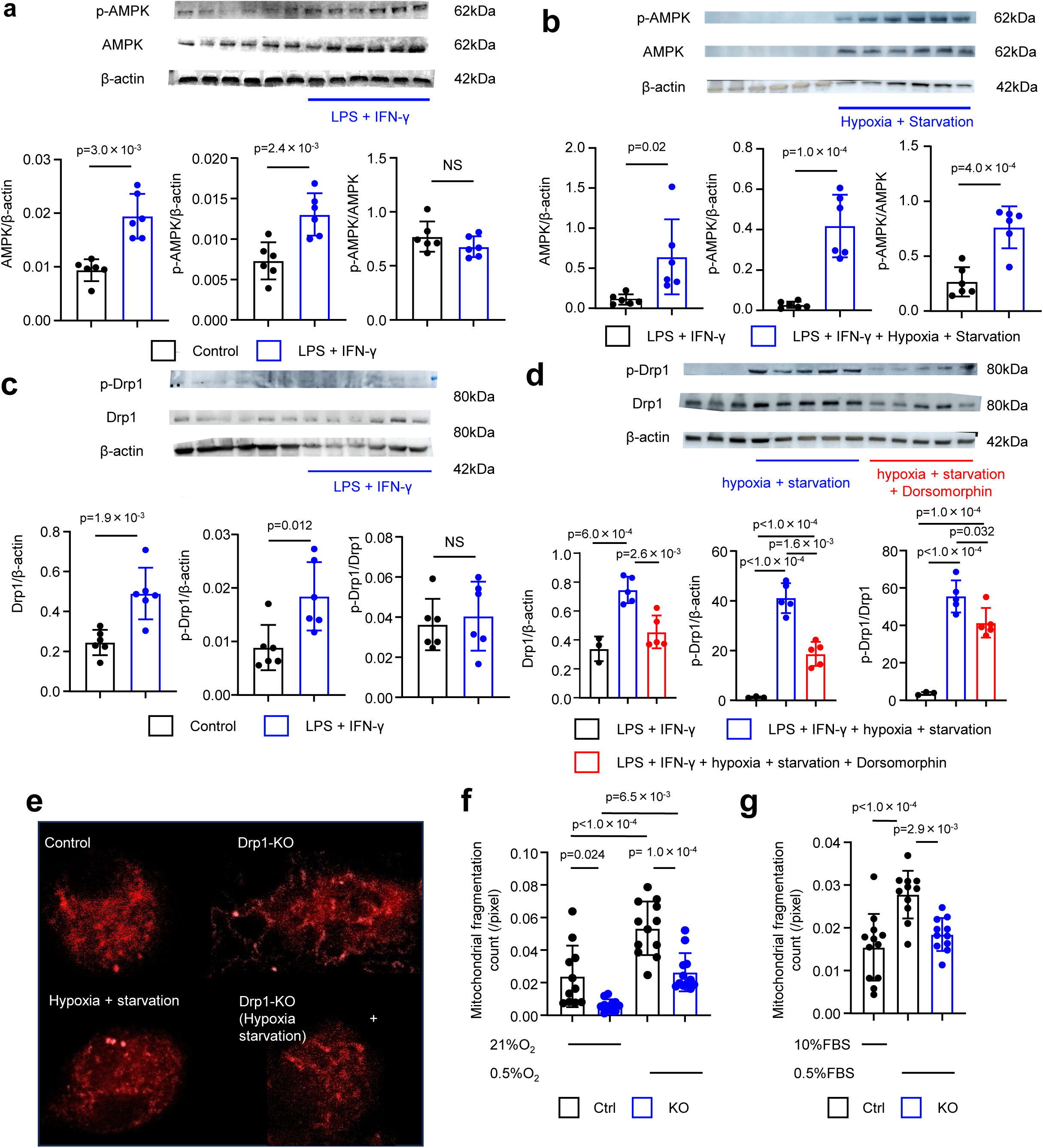
Hypoxia and starvation induce the phosphorylation of AMPK and subsequent activation of Drp1. (**a-d**) Western blot analysis and quantification of total AMPK, phosphorylated AMPK, total Drp1, phosphorylated Drp1, and β-actin in RAW264.7 cells at 24 hours after stimulation. LPS; 10 ng/ml, IFN-γ; 10 ng/ml, AMPK inhibitor (Dorsomorphin); 20 µM, hypoxia; 0.5% oxygen, starvation; 0.5% FBS. (**a**: p-AMPK/ AMPK; p=0.21, **c**: p-AMPK/ AMPK; p=0.64) (**a-c**; unpaired t-test) (**d**: one-way ANOVA with Bonferroni’s multi-comparison test) (**a**: N=6, **b**: N=6, **c**: N=6, **d**: N=3-5). (**e**) Mitochondria stained with MitoTracker. (**f-g**) mitochondrial fragmentation quantified after 24 hours of hypoxia (p=0.0029, ****p<0.0001 one-way ANOVA with Bonferroni’s multi-comparison test) (**f**: N=11-12), and starvation (**g**: N=11-12).). All data are presented as mean ± s.e.m. Ctrl: control, KO: Drp1-KO.

We previously reported that polarization toward inflammatory macrophages induces mitochondrial fission (*2*). However, it is unknown whether hypoxia or starvation induces morphological changes in the mitochondria of macrophages. Therefore, we have cultured thioglycolate-elicited peritoneal macrophages under hypoxia or serum-depleted conditions. As shown in Fig.3f, hypoxia induced mitochondrial fission, quantified as mitochondrial fragmentation count (MFC), and blockade of Drp1 by Mdivi-1 decreased MFC. Serum starvation also induced mitochondrial fission, and it was also decreased by Mdivi-1 (Fig.3g).

### Deletion of Drp1 decreased mitophagy in macrophages accumulating in infarcted myocardium

As the above data show, hypoxia and starvation, to which macrophages accumulating in the infarcted area are exposed, induced Drp1-mediated mitochondrial fission. It is, however, obscure how the deletion of Drp1 affects mitochondria in vivo. Hence, we observed macrophage mitochondria morphology in the infarcted tissue with transmission electron microscopy 3 days after MI (Fig.4a-d). In Extended Data Fig.2a, swollen mitochondria with destructive internal structures were regarded as damaged mitochondria. Extended Data Fig.2b demonstrates representative features of mitophagy, determined as autophagosomes or lysosomes containing mitochondria. There was an increase in mitochondrial area (2.39 vs 0.82×10^-2^µm^2^, p=1.0×10^-4^) and a decrease in mitophagy in the Drp1-KO group compared to the control group. (Fig.4e-g). The measurement of mitochondrial size by the length of the long and short axes indicated that the deletion of Drp1 increased the size of mitochondria compared to the control group (Fig.S4).

**Fig. 4.**
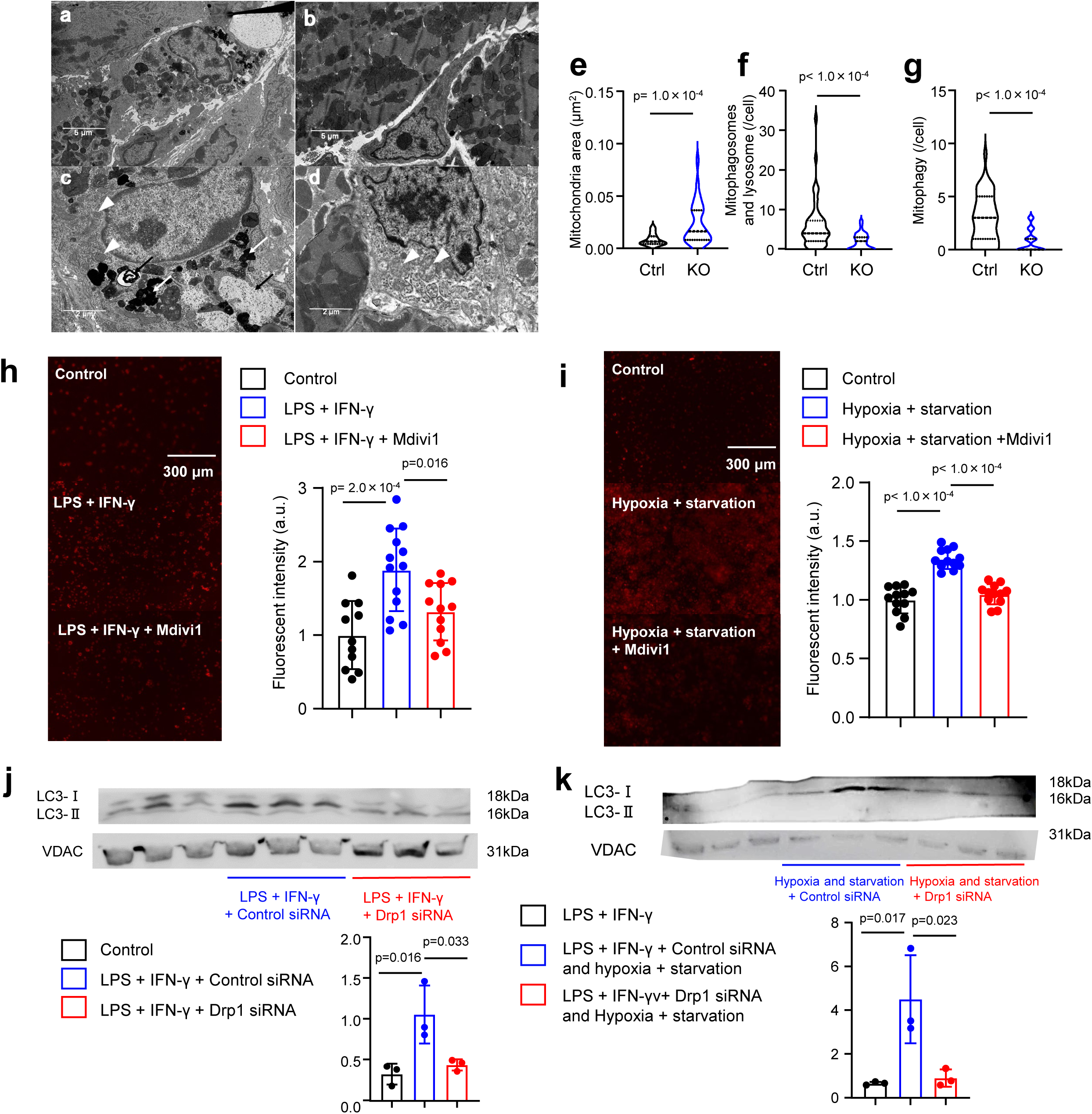
Deletion of Drp1 alleviates hypoxia- and starvation-induced mitophagy. (**a-b**) Macrophages accumulating in the infarcted myocardial tissues observed by transmission electron microscopy. a: control mouse, b: Drp1-KO mouse. (**c-d**) Magnified images of macrophages accumulating in the infarcted tissue. Mitochondria: white arrowheads, autophagosomes: white arrows, lysosomes: black arrowheads. c: control mouse, d: Drp1-KO mouse. (**e**) Size of mitochondria determined as an area in macrophages (unpaired t-test) (N=123 or 164). (**f**) Number of mitophagosomes and lysosomes per macrophage in each of the three mice (unpaired t-test) (N=49-50). (**g**) Number of mitophagy per macrophage in each of the three mice (unpaired t-test) (N=49-50). (**h-i**) Mitophagy is quantified using Mito-Keima Red in RAW264.7 cells treated with LPS, IFN-γ and/or 50 µM Mdivi-1 (one-way ANOVA with Bonferroni’s multi-comparison test) (N=11-13). (**j-k**) Western blot analysis of LC3 and VDAC in RAW264.7 cells treated with LPS, IFN-γ and/or 20 µM Drp1 siRNA (one-way ANOVA with Bonferroni’s multi-comparison test) (N=3). All data are presented as mean ± s.e.m. Ctrl: control, KO: Drp1-KO.

Then, mitophagy, a critical mechanism of mitochondrial quality control, was evaluated in a macrophage cell line, RAW264.7. In inflammatory macrophages induced by LPS and IFN-γ, mitophagy quantified by the fluorescence of mito-Keima Red was significantly suppressed by Drp1 inhibitor, Mdivi-1 (Fig.4h). Hypoxia and serum starvation-induced mitophagy was also suppressed by Mdivi-1 (Fig.4i). A Western blot analysis of LC3 using mitochondrial fraction of macrophages indicated that siRNA-mediated knockdown of Drp1 attenuated mitophagy that is underpinned by decreased LC3-II (Fig. 4j). Hypoxia and starvation-induced mitophagy were also attenuated by siRNA-mediated knockdown of Drp1 (Fig.4k). Mitophagy induced by CCCP and oligomycin, which is a commonly used method in vitro, was also decreased by Drp1 siRNA, as evidenced by Western blot analysis and mito-Keima Red (Fig.S3).

### Macrophage transcriptome analysis demonstrated that long-term inhibition of Drp1 induces the expression of genes associated with inflammation

RNA-sequencing was conducted in RAW264.7 cells treated with LPS and IFN-γ to evaluate the effect of Drp1 inhibition on the transcriptome of macrophages, especially focusing on the duration of Drp1 inhibition─early phase (24 hours) vs. late phase (96 hours) (Fig.5a-b). Principal component analysis indicated distinct gene expression patterns between control and Mdivi-1-treated RAW264.7 cells (Fig.5c-d). The scatter plot indicates upregulated and downregulated genes at each time point. In the early phase. i.e., 24 hours after treatment, genes including *Fn1*, *Thbs1*, *Cola12*, *Col1a1*, *Sparc*, *Serpinh1*, *Fpr1*, *Fbin2*, *Cyp1b* and *Heg1* were decreased by blockade of Drp1 (Fig.5e). GO analysis demonstrated that differentially expressed genes (DEGs) include genes associated with nuclear division and cell cycle regulation (Extended Data Fig.4a). In the later phase, i.e., 96 hours after treatment, genes including *mt-Co3*, *Fth1*, *Il1b*, *Eef1a1*, *Il1a*, *Ptgs2*, *Hspa8*, *Ubc* and *Ccl3* were increased by blockade of Drp1 (Fig.5f). GO analysis and enrichment analysis in transcription factor targets indicated changes of biological process and transcription factors at this point (Extended Data Fig.5c-f).

Network analysis suggested that cell cycle-related molecules, particularly those involved in mitosis, such as Kif11 and Bub1b, may function as hub genes among the DEGs 24 hours after Mdivi-1 treatment (Fig.5g). In contrast, hub genes after 96 hours of Mdivi-1 treatment included genes associated with inflammation, e.g., *Il1b* and *Il6* (Fig.5h). GSEA analysis indicates that inhibition of Drp1 for 96 hours suppresses expression of genes associated with inflammation, especially polarity shift towards inflammatory phenotype (Fig.5i).

### Long-term inhibition of Drp1 impairs mitochondrial quality control machinery and causes sustained inflammation in macrophages

To investigate the detailed molecular mechanisms of how inhibition of Drp1 modifies inflammation in macrophages, we assessed the mitochondrial DNA (mtDNA) leakage from mitochondria to the cytosol. In RAW264.7 cells treated with the Drp1 inhibitor Mdivi-1, DNA was extracted from the cytosolic fraction, and mtDNA was quantified by semi-quantitative real-time PCR using primers for cytochrome c oxidase I, a molecule encoded by mtDNA. Treatment with Mdivi-1 for 24 hours did not significantly change the cytosolic mtDNA amount compared to control RAW264.7. However, autophagy inducer Tat-beclin1 significantly decreased cytosolic mtDNA compared to macrophages treated with Mdiiv-1 (Fig.6a1). In contrast, Mdivi-1 treatment for 96 hours increased cytosolic mtDNA, which was mitigated by a voltage-dependent anion channel (VDAC) oligomerization inhibitor VBIT-4 or Tat-beclin1 either (Fig.6a2). Hypoxia and starvation increased cytosolic mtDNA at an earlier time point, 24 hours, which was also inhibited by either VBIT-4 or Tat-beclin1 (Fig.6a3).

Then, inflammatory cytokine expressions were evaluated. In these macrophages, Mdivi-1 partially reduced the mRNA expression of *Tnf* and *Il6* after 24 hours of treatment (Fig.6b). An autophagy inducer, Tat-beclin1 decreased mRNA expression of *Il1b*, *Ccl2*, *Tnf*, and *Il6* compared to macrophages treated with LPS and IFN-γ at this time point. Expressions of *Il1b*, *Ccl2*, *Tnf*, and *Il6* were increased after 96 hours of Mdivi-1 treatment, all of which were attenuated by Tat-beclin1 (Fig.6c). Under conditions with hypoxia and starvation, Mdivi-1 induced expression of *Il1b*, *Tnf*, and *Il6* at 24 hours after treatment, and which was abrogated by Tat-beclin1. These results suggest that inhibition of Drp1 suppresses the expression of inflammatory cytokines at the early time point. In contrast, the inhibition of Drp1 for a long time increases the expression of these inflammatory cytokines, possibly via the impairment of mitochondrial quality control mechanisms, including mitophagy. Under conditions with hypoxia and energy deprivation, inhibition of Drp1 accelerated the induction of inflammatory processes.

### Macrophage ZBP1 plays essential roles in the exacerbation of inflammation and post-infarct LV remodeling after blockade of Drp1

The above results suggest that blockade of Drp1 impairs mitophagy and induces leakage of mtDNA from damaged mitochondria to the cytosol, which could induce sterile inflammation. The role of an intracellular nuclear acid receptor was examined to address the detailed molecular mechanisms by which mtDNA induces inflammation. In the cytosol, there are several nuclear receptors, but we focused on Z-DNA Binding Protein 1 (ZBP1) because recent evidence suggests that ZBP1 is a DNA sensor recognizing mtDNA released from cells exposed to oxidative stress (*8*). To clarify the specific role of ZBP1 in myeloid cells, including macrophages, bone marrow transplantation was performed using ZBP1-KO mice as donors (BM-ZBP1-KO). Control mice transplanted bone marrow cells from wild-type mice were prepared as controls (BM-Control). In Myeloid ZBP1-KO mice, inhibition of Drp1 by Mdivi-1 significantly decreased LVEF and increased LV dimensions, as observed in Drp1-KO mice. In these mice, genetic deletion of ZBP1 in myeloid cells partially attenuated the reduction of LVEF (Fig.7a). Enlargement of LV dimensions was abrogated in BM-ZBP1-KO mice compared to BM-Control treated with Mdivi-1 (Fig.7b-c). Deletion of ZBP1 in myeloid cells decreased the circumferential length of infarcted tissue and reduced collagen content, as shown in Fig.7d-f. Macrophage accumulation in and around the infarcted myocardial tissue was attenuated in BM-ZBP1-KO mice (Fig.7f). Moreover, immunohistochemical analysis of cleaved caspase-3 at day 28 after MI indicated that deletion of macrophage ZBP1 rather decreased the ratio of cleaved caspase-3^+^ apoptotic cells in the peri-infarct border area (Fig.7g).

In in vitro cultured RAW264.7 cells in which polarization toward inflammatory macrophages was induced, Mdivi-1 induced apoptotic cell death 24 hours after the treatments identified as TUNEL^+^ cells. siRNA-mediated knockdown of ZBP1 abrogated the induction of apoptotic cell death by Mdivi-1 (Fig.7h). In term of inflammatory cytokines, induction of *Il1b*, *Ccl2*, *Tnf*, and *Il6* mRNAs at 96 hours after Mdivi-1 treatment was abrogated by siRNA-mediated knockdown of ZBP1 (Fig.7i). Hypoxia and starvation in these RAW264.7 cells did not significantly increased mRNA expressions of *Il1b*, *Tnf*, and *Il6*. However, the knockdown of ZBP-1 suppressed the expression of these molecules until a comparable level to control RAW264.7 cells cultured in normoxic conditions without Mdivi-1 (Fig.7j).

## Discussion

Here, we have clarified the specific role of macrophage Drp1 in the healing process of infarcted tissues in the heart, especially focusing on the regulatory mechanisms of macrophage-mediated inflammation. Novel findings include that macrophage-selective deletion of Drp1 exacerbated inflammation and LV remodeling after MI. In addition, in inflammatory macrophages especially exposed to interruption of oxygen and energy supply, AMPK phosphorylation induced downstream Drp1 activation and mitochondrial fission. In macrophages accumulating in infarcted tissues, mitophagy is impaired by the deletion of Drp1, which induces cytosolic leakage of mtDNA. These mtDNAs induced sterile inflammation via ZBP-1-mediated mechanisms, which is underpinned by the in vivo observation that myeloid-specific deletion of ZBP-1 abrogated exacerbation of LV remodeling by blockade of Drp1.

Mitochondria, intracellular organelles existing in most eukaryotic cells, play central roles in cellular energy production. ATP is synthesized in mitochondria through glycolysis, the tricarboxylic acid (TCA) cycle, and the electron transport chain. Mitochondria also play essential roles in the production of reactive oxygen species, regulation of cell death, and calcium storage (*9*). mtDNA, transmitted by maternal inheritance and has different features from genomic DNA, is involved in various biological processes (*10*). Mitochondria continuously change their morphology under physiological and pathological conditions, which are called mitochondrial dynamics (*2–7, 9*). By specifically intervening with the mitochondrial pro-fission molecule Drp1 in monocytes/macrophages, we have elucidated the role of mitochondrial fission in monocytes/macrophages, especially in monocytes/macrophase-mediated inflammation after myocardial infarction.

In the context of monocytes/macrophages, there are two distinct phases after MI. In the early phase, inflammatory monocytes/macrophages accumulate in the infarcted area, peaking on days 3-4 after MI (*11*). Subsequently, reparative macrophages become the dominant population. Deletion of Drp1 in macrophages caused sustained macrophage accumulation in the infarcted heart until 28 days after MI. Flow cytometry indicated that deletion of Drp1 causes sustained accumulation of the Ly-6C^high^ subset, known as inflammatory monocytes, which in turn resulted in the reduction of the absolute number of cells included in the Ly-6C^low^ subset containing reparative macrophages. The decrease in monocytes in the bone marrow and the spleen suggests that the mobilization of monocytes was accelerated in Drp1-deficient mice. The survival curve demonstrated that deaths after MI frequently occurred until 10 days after LAD ligation, suggesting that the cause of death was a cardiac rupture in most of the cases due to impairment of the healing process and mechanical fragility of the infarcted tissues. Postmortal autopsy indicated that heart failure was a minor cause of death during the follow-up period. Pathological findings suggested that ballooning of the infarcted tissues, promoted by sustained inflammation in the infarcted and peri-infarcted area, was the primary mechanism of LV remodeling rather than compensated cardiac hypertrophy in non-infarcted myocardium.

Transmission electron microscopy revealed that deletion of Drp1 in macrophages increased not only the length of mitochondria but also the mitochondrial volume, suggesting accumulation of damaged mitochondria in Drp1-deficient macrophages. Mitophagy usually removes damaged mitochondria from cytoplasms, but mitophagosomes were decreased in Drp1-deficient macrophages. It is speculated that Drp1-mediated mitochondrial fission separates damaged parts of mitochondria from healthy ones. However, the damage of mitochondria beyond a certain threshold, mitochondria is eliminated by mitophagy. In this process, the PINK1-Parkin-LC3-mediated mechanism is well-known (*4–6, 9, 12*). Therefore, we evaluated mitophagy by immunoblotting LC3 in addition to quantification with mitophagy indicator mito-Keima. The finding that mitophagy is impaired in Drp1-deficient macrophages suggests that a reduction of mitochondrial size through mitochondrial fission promotes the incorporation of damaged mitochondria into mitophagosomes, although other reports are suggesting the existence of mitophagy irrespective of mitochondrial size (*13*). It is speculated that Drp1 promotes mitophagy through these mechanisms in macrophages.

After MI, circulating leukocytes, particularly inflammatory monocytes, accumulate in the heart, notably within the infarcted tissues and its penumbra. In the infarcted tissues, where the blood supply is disrupted, accumulated macrophages are exposed to hypoxia and a shortage of energy sources. Under these conditions, it is speculated that AMPK, a well-known energy sensor activated by ATP/ADP imbalance, is activated. Therefore, we have quantified AMPK phosphorylation in inflammatory macrophages and confirmed that hypoxia and starvation increased the phosphorylation of AMPK. The results showing that an AMPK inhibitor partially inhibited hypoxia and starvation-induced Drp1 activation determined by phosphorylation at Ser616 indicate that AMPK could be an upstream activator of Drp1 in macrophages accumulating to the infarcted tissues (*14*). Along with activation of Drp1, mitochondrial fissions were induced by hypoxia and starvation in macrophages, whereas it was attenuated in macrophages from macrophage Drp1-deficient mice. These findings align with our previous research, which demonstrated that mitochondria in inflammatory macrophages are more fragmented than non-stimulated ones (*2*).

Transcriptome analysis of macrophages indicated that blockade of Drp1 induces distinct gene expression between 24 hours and 96 hours after treatment with Mdivi-1 (Fig.5). In the early phase, inhibition of Drp1 induced the expression changes of genes associated with mitosis. This result suggests not only that mitochondrial fission occurs during mitosis but also that inhibition of mitochondrial fission impairs cell proliferation. After long-term inhibition of Drp1, genes associated with inflammation and catabolic processes were induced. Therefore, we have focused on molecular mechanisms by which the accumulation of damaged mitochondria triggers inflammation in macrophages. We particularly focused on mtDNA as a key factor. As demonstrated in Figure 6a, the blockade of Drp1 for 96 hours, but not for 24 hours, increased cytosolic mtDNA in inflammatory macrophages. In macrophages exposed to hypoxia and starvation, mtDNA was increased in the cytosol after 24 hours of treatment with Mdivi-1. The results indicating that VBIT-4, a small-molecule inhibitor of VDAC, abrogated the leakage of mtDNA suggest that mitochondrial outer membrane pores (MOMP) formed by VDAC oligomerization are involved in the process of mtDNA leakage from the mitochondrial matrix to the cytosol. Moreover, these findings are supported by Kim’s report, which states that during mitochondrial elongation, mtDNA is released through MOMP, requiring the opening of the mitochondrial permeability transition pore for inner membrane permeabilization (*10*). The data that Tat-beclin 1, an inducer of autophagy, also abrogated the leakage of mtDNA indicates that impaired mitophagy could be one of the mechanisms that cause the accumulation of damaged mitochondria and resultant mtRNA leakages. The expression of inflammatory cytokines was paralleled with the results of cytosolic mtDNA. Further induction of inflammatory gene expressions by blockade of Drp1 was not observed in inflammatory macrophages treated with Mdivi-1 for 24 hours, whereas blockade of Drp1 for 96 hours further increased mRNA expression of *Il1b*, *Ccl2*, *Tnf*, and *Il6*. These inductions were similarly abrogated by Tat-beclin 1. It was also parallel to the mtDNA that hypoxia and starvation shorten the time until the inflammatory cytokines are induced. These results suggest that while Drp1 deletion alleviates acute inflammation, it induces sustained inflammation in the later phase.

**Fig. 5.**
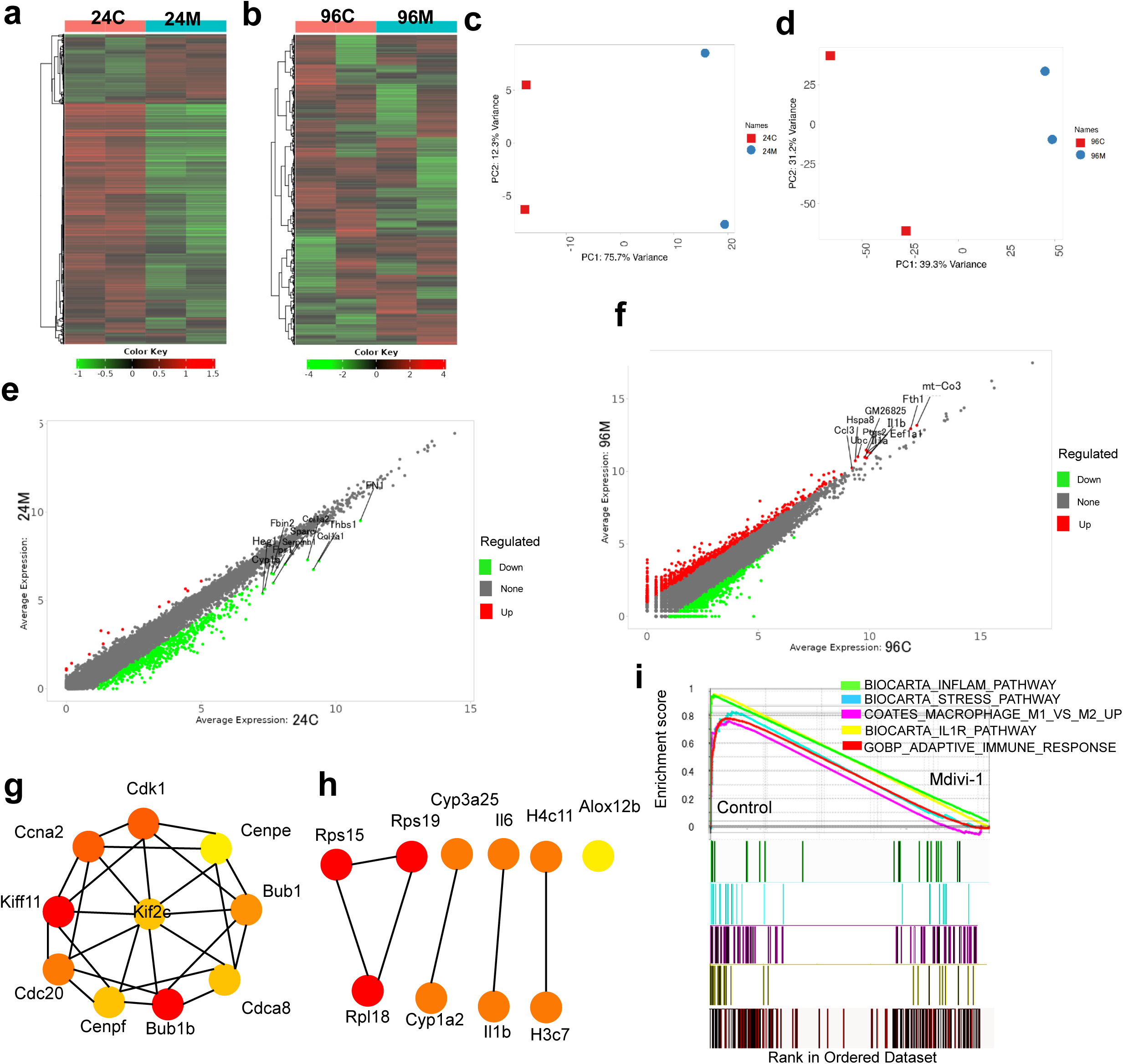
Inhibition of Drp1 for 24 hours and 96 hours induces distinct gene expressions in macrophage. (**a-b**) Heatmap of top 500 differentially expressed genes in RAW264.7 cells. 24C and 96C: LPS and IFN-γ for 24 or 96 hours, 24M and 96M: LPS, IFN-γ, and Mdivi-1 for 24 or 96 hours, respectively. (**c-d**) Principal component analysis at 24 hours (c) or 96 hours (d) after treatment. (**e-f**) Scatter plot of genes after 24-hour (e) or 96-hour (f) treatment. (**g-h**) The top 10 hub genes were analyzed in 24 hours (g) or 96 hours (h) treated samples using Cytoscape version 3.10.2 and cytoHubba 0.1. (g) or 96 hours (h) after treatment. The red color indicates top-scored genes. (**i**) GSEA analysis of RAW264.7 cells treated with or without Mdivi-1 for 96 hours. All the RNA-seq analyses were repeated twice using different series of samples.

**Fig. 6.**
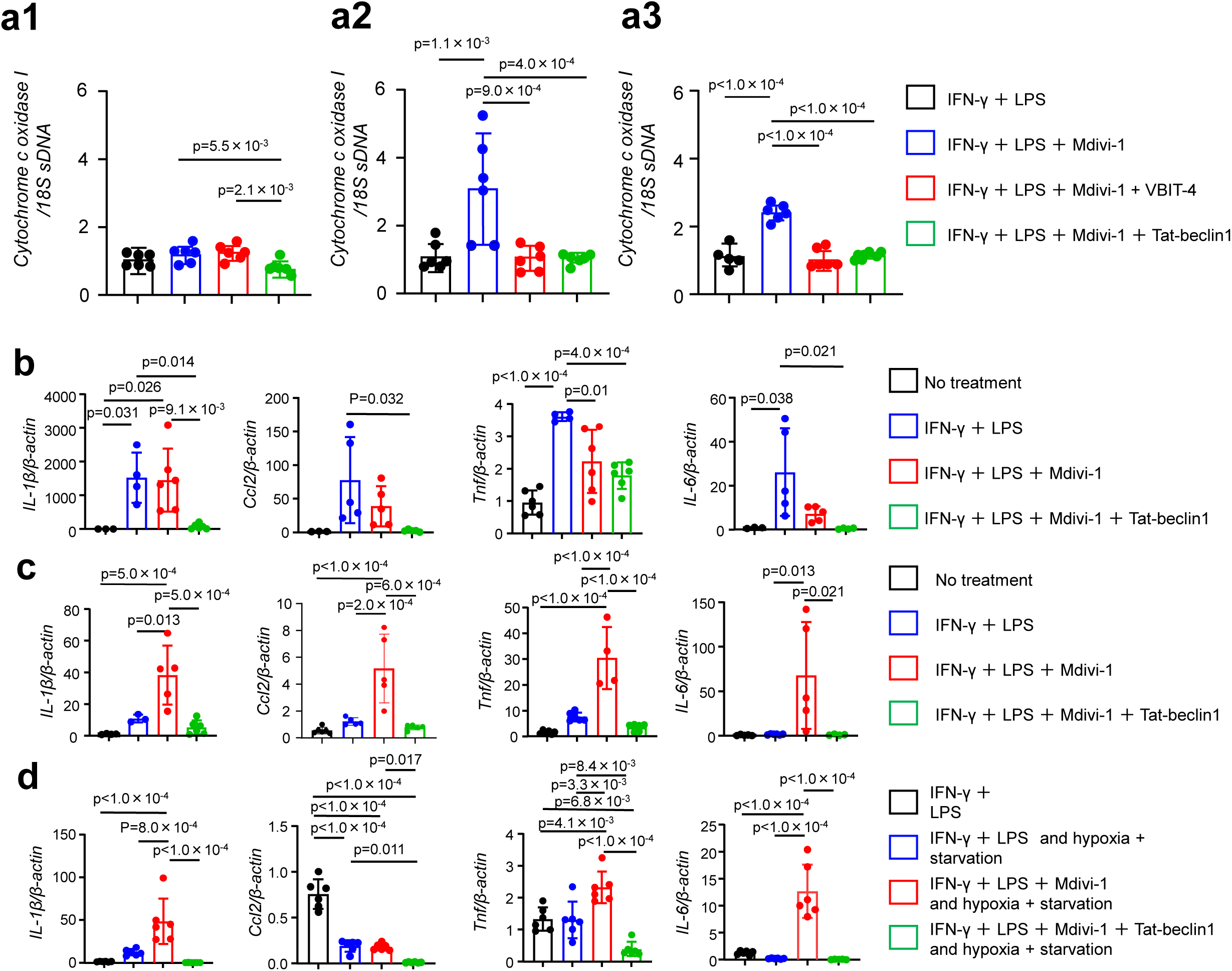
Impairment of mitophagy causes cytosolic mitochondrial DNA release and sustained inflammation. (**a**) Cytochrome c oxidase I DNA in cytosolic fraction was quantified by semi-quantitative real-time PCR at 24 hours (a1), 96 hours (a2) after treatment, or 24 hours after treatment under hypoxia and starvation (a3). (one-way ANOVA with Bonferroni’s multi-comparison test) (N=5-6). VBIT-4; 20 µM, Tat-beclin1; 25 µM. (**b-c**) The mRNA expression of inflammatory cytokines at 24 hours (b) and 96 hours (c) after treatments (one-way ANOVA with Bonferroni’s multi-comparison test) (N=4∼6). (**d**) The mRNA expression of inflammatory cytokines at 24 hours after treatments under hypoxia and starvation (one-way ANOVA with Bonferroni’s multi-comparison test) (N=4∼6). All data are presented as mean ± s.e.m.

The polarity shift to reparative macrophages is also crucial in the process of tissue repair after MI to promote the healing and scarring of the infarcted tissue. Therefore, we have investigated the effect of Drp1 inhibition on the polarity shift toward less inflammatory reparative macrophages. The data showed that inhibition of Drp1 induced the expression of molecules associated with reparative macrophages, including CD206 and Arg1, and these changes were abolished in hypoxia and starvation conditions (Fig.S1c). Moreover, exposure to hypoxia and starvation likely shifts M2 macrophages to an M1 phenotype when deletion of Drp1 does not affect the allocation of macrophages, although it leads to a shift of macrophages from M1 to M2 phenotype. Previous research indicates that mitochondrial dysfunction inhibits the repolarization of M1 to M2 macrophages (*15*). This result suggests that the inflammatory milieu in the infarcted tissue could skew the M2 macrophages towards an M1 phenotype as a potential mechanism of sustained inflammation.

What are the downstream pathways of mtDNA? Recent reports have identified several cytosolic DNA sensors, including Toll-like receptor 9 (TLR-9), cGAS, STING, Absent in Melanoma 2 (AIM2), and IFN-γ-inducible 16 (IFI16) (*16*). Additionally, numerous studies have highlighted that ZBP1 acts as a cytosolic nucleic acid sensor for components such as mtDNA. It not only induces apoptosis through RIPK3 but also activates inflammatory signaling pathways mediated by non-degradative ubiquitin chains, utilizing Receptor-Interacting Protein Kinase 1 (RIPK1) and RIPK3 as scaffolds (*9–10, 17–21*). Our data indicates that ZBP1 is a critical mediator of mtDNA-induced inflammation, contributing to LV remodeling. According to a recent study, ZBP1 acts as an innate immune sensor of the mitochondrial genome, amplifying cGAS-STING-dependent Type I interferon (IFN-I) signaling in both human and mouse cells (*10*). Another study revealed that cGAS-STING and RIPK3 are downstream pathways influenced by ZBP1 (*18*). Figures 7i and 7j further reveal that the elevated cytokine levels during the subacute and acute phases under hypoxia and starvation were suppressed by Tat-Beclin1 and siRNA-mediated knockdown of ZBP1. These findings implicate ZBP1 in the downstream signaling pathways modulated by Drp1. Damaged mitochondria that are not cleared by mitophagy may release mtDNA, thereby contributing to inflammation via ZBP1. Over 96 hours, there was an increase in *Il1b*, *Ccl2*, *Tnf*, and *Il6* mRNAs, whereas, at 24 hours under hypoxia and starvation, there was an increase in *Il1b*, *Tnf*, and *Il6* without *Ccl2*. Figures 7a through 7e show that macrophage ZBP1-KO improved LV remodeling, including cardiac fibrosis exacerbated by blockade of Drp1. Figure 7f revealed a significant decrease in macrophage accumulation in the infarcted hearts of ZBP1-KO mice compared to control mice treated with Mdivi-1. Figures 7g and 7h confirmed that ZBP1-KO alleviated apoptosis in macrophages post-MI. This study indicates that ZBP1 activation by mtDNA released from damaged or dysfunctional mitochondria can exacerbate inflammation, leading to LV remodeling.

**Fig. 7.**
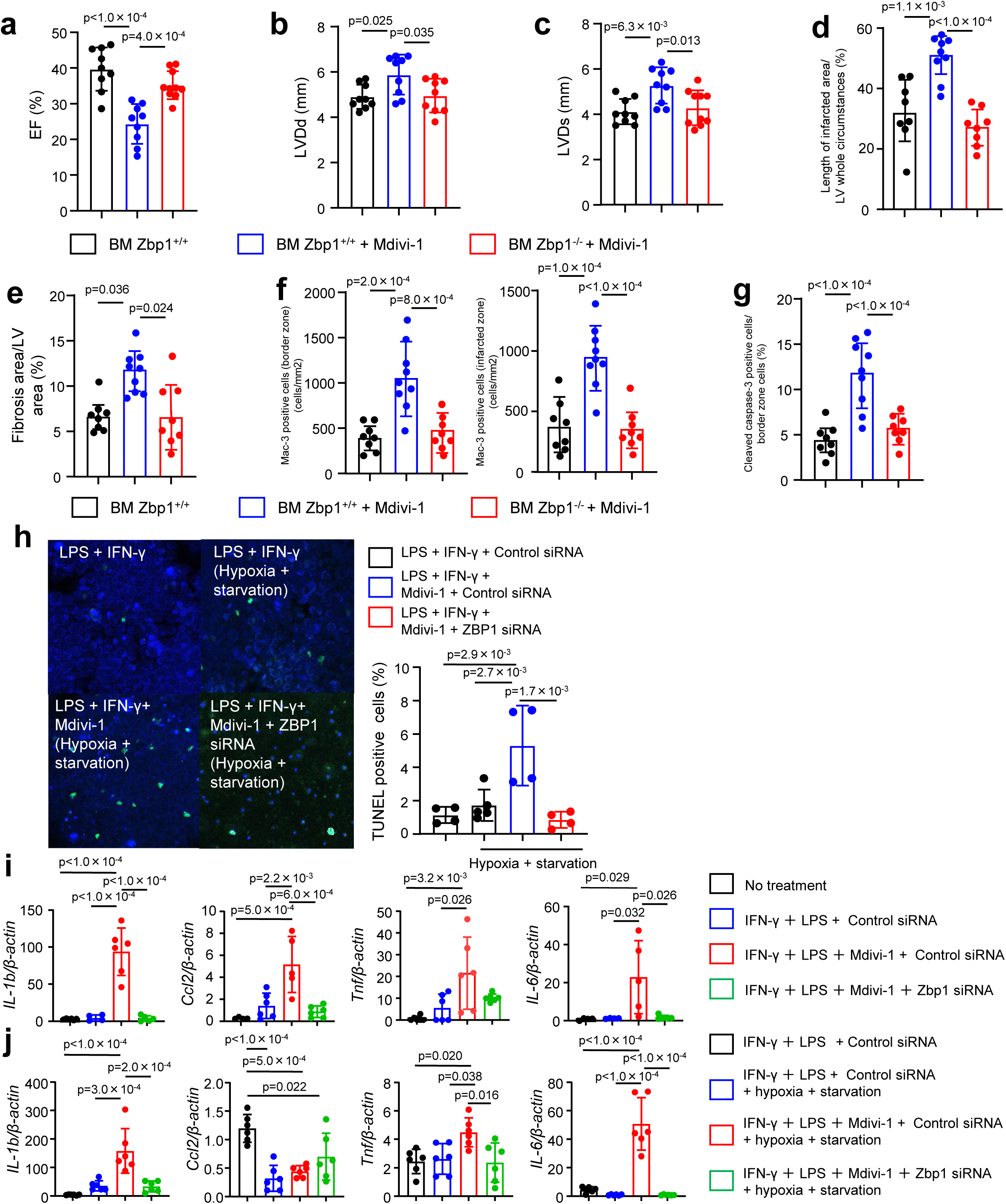
Deletion of ZBP1 in myeloid cells abrogated exacerbation of LV remodeling by blockade of Drp1. (**a-c**) The data of echocardiography in control (BM Zbp1^+/+^) and myeloid-specific ZBP1-KO (BM Zbp1^-/-^). (one-way ANOVA with Bonferroni’s multi-comparison test) (N=9-10). (**d**) The relative length of the infarcted area to the LV circumstances (student’s t-test) (N=8-9). (**e**) Quantitative results of fibrosis relative to the total LV area (one-way ANOVA with Bonferroni’s multi-comparison test)(N=8-9). (**f**) Accumulation of mac-3 positive cells in the border zone and infarcted area of the heart (one-way ANOVA with Bonferroni’s multi-comparison test)(N=8-9). (**g**) The rate of apoptotic cells relative to total cells in the border areas (one-way ANOVA with Bonferroni’s multi-comparison test) (N=8-9). (**h**) TUNEL staining in RAW264.7 cells (left) and quantitative results (right) (one-way ANOVA with Bonferroni’s multi-comparison test) (N=4-5). (**i-j**) The mRNA expression of inflammatory cytokines determined by real-time PCR at 96 hours after treatments. (one-way ANOVA with Bonferroni’s multi-comparison test) (N=4 to 6). All data are presented as mean ± s.e.m. BM: Bone marrow

In conclusion, this study has clearly shown the critical role of macrophage Drp1 in inflammation and LV remodeling after MI. Drp1-mediated mitochondrial fission plays critical roles in mitochondrial quality control mechanisms and could be a novel therapeutic target for the prevention of chronic heart failure after MI.

## Acknowledgments

We thank Tomoko Fujita and Kozue Nakamura for their excellent technical assistance. We also appreciate the Shared-Use Research Center (SRC) of the University of Occupational and Environmental Health, Japan, especially Dr. Mitsuru Yokoyama, for excellent technical assistance in electron microscopy imaging.

## Sources of Funding

Ministry of Education, Culture, Sports, Science, and Technology (MEXT) (Grants-in-Aid for Scientific Research 17K16011, 19K08518 to J.K.)

University of Occupational and Environmental Health (UOEH Grant-in-Aid for Young Researchers to J.K.).

## Disclosures

None

## Novelty and Significance

### What is known?

- Macrophages, consisting of various subsets, from inflammatory to reparative ones, play a central role in modulating the healing process after MI.
- Mitochondrial dynamics, i.e., morphological changes of mitochondria, are critical in some cardiovascular diseases.
- Inhibition of Drp1 in macrophages has been shown to reduce inflammation in vascular injury models.

### What new information does this article contribute?

- Under hypoxia and starvation, conditions similar to infarcted areas after MI, AMPK is activated, leading to Drp1 activation in macrophages.
- In macrophages with Drp1 deficiency, the accumulation of damaged mitochondria due to impaired mitophagy causes mtDNA leakage into the cytoplasm, leading to sterile inflammation via the ZBP1 pathway.
- Deletion of macrophage Drp1 significantly increased mortality due to LV rupture in the early phase and exacerbated LV remodeling in the later phase after MI.

## Notes

### Competing Interest Statement

The authors have declared no competing interest.

